# A Complexity-Science Framework for Studying Flow: Using Media to Probe Brain-Phenomenology Dynamics

**DOI:** 10.1101/2025.07.11.664430

**Authors:** Fran Hancock, Rachael Kee, Fernando Rosas, Manesh Girn, Steven Kotler, Michael Mannino, Richard Huskey

## Abstract

Consciousness spans a range of phenomenological experiences, from effortless immersion to disengaged monotony, yet how such phenomenology emerges from brain activity is not well understood. Flow, a phenomenological experience frequently elicited by interactive media, has drawn attention for its links to performance and wellbeing, but existing neural accounts rely on single region or small network analyses that overlook the brain’s distributed and dynamic nature. Complexity science offers tools that capture brain-wide dynamics, but this approach has rarely been applied to flow or to its natural comparisons: boredom and frustration. Consequently, it remains unclear whether tools drawn from complexity science can objectively discriminate between these phenomenological experiences while also clarifying their neural basis. To address this uncertainty, we induced each phenomenological experience with a difficulty-titrated video game during functional magnetic resonance imaging and collected concurrent behavioral and self-report data. Our complex systems analyses revealed that flow, in this experimental setup, shows an inverse relationship to global entropy with moderate explanatory power, and is not explained by either synchronization or metastability, whereas boredom and frustration exhibit different configurations of brain-dynamics metrics. Notably, these findings integrate previously separate prefrontal and network-synchrony observations within a single dynamical systems framework and identify complexity-based markers with the potential to map the neural underpinnings of media-related benefits.

## 1. Introduction

Human consciousness encompasses an array of phenomenological experiences, each representing a dynamic interplay of perception, cognition, and emotion. The modern scientific study of phenomenology was pioneered in the late 19th century by William James (1890), who argued that emotional experiences are characterized by dynamic interactions between our internal self and the external world. In the years since, researchers have investigated an ever-growing variety of phenomenological experiences that characterize typical and abnormal brain function. Such inquiry still includes topics such as affect and emotion (Barrett, 2017; Cacioppo et al., 1999), but has since expanded to encompass additional experiences, including meditation (Van Dam et al., 2018), mindfulness (Zhou et al., 2023), self-transcendence (Kang et al., 2018), self-affirmation (Falk et al., 2015), acute psychosis (Horga & Abi-Dargham, 2019; Insel, 2010), sleep and dreaming (Solms, 2021), flow (Csikszentmihalyi, 1990), the psychedelic experience (Carhart-Harris et al., 2014; Carhart-Harris & Friston, 2019; Girn et al., 2023), audience responses to media (Grall et al., 2021; Hasson et al., 2008; Schmälzle, 2022; Schmälzle et al., 2024; Schmälzle & Grall, 2020; Schmälzle & Huskey, 2023a; Weber et al., 2024), player responses to video games (K. Mathiak & Weber, 2006; Weber et al., 2006, 2018), and more. Descriptive accounts of these phenomenological experiences abound, as do proposed mechanisms. Yet, a conclusive understanding of their neural substrates remains elusive. Why?

A significant challenge lies in the difficulty of studying these experiences within controlled experimental paradigms. Advances in naturalistic neuroimaging (Finn et al., 2022; K. Mathiak & Weber, 2006; Schmälzle & Huskey, 2023a; Sonkusare et al., 2019) have addressed some limitations, but the majority of cognitive neuroscience often remains confined to models focused on isolated cognitive processes (e.g., attentional vigilance, working memory, reward signaling) within a single (i.e., locationist) or relatively small set (i.e., connectionist) of brain regions (Noble et al., 2024). This reductionist approach persists despite evidence that large-scale, whole-brain dynamics underlie a broad spectrum of cognitive functions (Westlin et al., 2023), including relatively low-level processes such as strategic attention and recognition memory (Davison et al., 2015; Telesford et al., 2016). If lower-level cognitive processes resist simple locationist or connectionist models, then what hope is there for using these paradigms to explain complex phenomenology implicating multiple interacting cognitive processes?

Neurophenomenology (Lutz & Thompson, 2003) connects first-person experience with moment-by-moment brain dynamics. Complexity science extends this bridge by modeling the brain as a nonlinear dynamical system and quantifying its emergent, information-rich patterns; thereby providing insights that surpass region-by-region accounts (Bassett & Gazzaniga, 2011; Hancock, Rosas, et al., 2022; Kelso, 1995; Seguin et al., 2023). In this article, we focus on flow experiences (Csikszentmihalyi, 1990) as a test-case for studying complex phenomenological experiences from a complexity science perspective. We use media to elicit flow, which allows us to observe linkages between neural complexity signatures and media-elicited phenomenological experience in a way that clarifies longstanding questions in flow research and provides a roadmap for inquiry into the neural basis of a wide variety of phenomenological experiences, including those elicited by media, and their subsequent outcomes (Weber et al., 2015).

### 1.1 Flow, Media, and Wellbeing

Flow, first described five decades ago by psychologist Mihaly Csikszentmihalyi (1975), is an emergent phenomenological experience characterized by focused attention, altered temporal awareness, reduced self-reflective thought, and high intrinsic reward. Flow is implicated in a variety of positive wellbeing outcomes, including successful goal pursuit (Bernecker & Becker, 2021; Tepper & Lewis, 2021), increased resilience (Tabibnia, 2020), adaptive coping during uncertainty (Rankin et al., 2019; Sweeny et al., 2020), resistance to burnout (Mosing et al., 2018), and more (Csikszentmihalyi, 1990). Importantly, media can elicit flow (Sherry, 2004), and experiencing flow during media use has also been linked to a number of wellbeing outcomes (Chiang et al., 2011; Vella et al., 2013).

Despite years of research, the neural mechanisms underlying flow remain fragmented, with competing models (for a review, see Kee & Huskey, in press). One influential model posits that flow results from a large-scale deactivation of the prefrontal cortex (Dietrich, 2004), whereas another argues that medial prefrontal cortex downregulation, causally driven by the dorsal raphe nucleus, contributes to flow (Ulrich et al., 2016a).^1^ Flow has also been theorized as resulting from a synchronization between attentional and reward networks (Weber et al., 2009).^2^ Finally, another model argues that these earlier specifications can be integrated by considering the cerebellum (Gold & Ciorciari, 2021).

These models, often developed from different experimental protocols, data analysis strategies, theoretical and philosophical commitments, have struggled to converge on a unified account of flow’s dynamic neural basis. A source of this lack of consensus could be the limitations of reductionist frameworks that focus on isolated regions (locationist) or static networks (connectionist), rather than the distributed, time-varying patterns that characterize complex phenomenological experiences.

### 1.2 The Present Study

Building on recent work (Girn et al., 2023; Pessoa, 2022; Turkheimer et al., 2022), we argue that flow is best understood using the latest tools and perspectives of complexity science (Hancock, Rosas, et al., 2022; Hancock et al., 2025; Turkheimer et al., 2022). Complexity science seeks to uncover general laws governing complex systems, integrating concepts from statistical physics, dynamical systems theory, and information theory (Holland, 2014; Thurner et al., 2018; Waldrop, 1993). It conceptualizes the brain as a complex system composed of discrete but integrated subunits that interact in non-linear ways to give rise to spontaneously organizing macroscopic patterns that cannot be reduced to the individual parts; that is, a complex system with the properties of multiplicity, interdependence, non-linearity, self-organization, and emergence (Jensen, 1998; Ladyman et al., 2013).

In this study, we manipulated the phenomenological experience of flow—using boredom and frustration as canonical comparisons as well as rest—and used tools drawn from complexity science to challenge and potentially unify pre-existing neuroscientific models. We then applied a comprehensive set of complexity analyses to functional magnetic resonance imaging (fMRI) data and compared them across conditions. Subsequently, we examined relationships with the subjective experiences of flow and enjoyment, as well as with a behavioral correlate of flow. Our objective was to understand which, if any, of these complexity signatures is most sensitive to differences between flow, boredom, and frustration, as well as which best explain subjective and behavioral measures. In doing so, our study directly advances this special issue’s goal of investigating the neural mechanisms that mediate the risks and benefits of modern media use. Moreover, we identify complexity-based signatures of coordination dynamics that guide future confirmatory research regarding the neural underpinnings of flow.

## 2. Methods

### 2.1 Previous Reporting and General Overview

The fMRI and behavioral data analyzed in this manuscript were previously reported on by Huskey and colleagues (2022). This earlier experiment successfully manipulated flow, boredom, and frustration (as confirmed via self-report), and evaluated fMRI data from a network neuroscience perspective (Fornito et al., 2016; Sporns, 2011). Here we completely re-evaluate the data, this time from a complexity science perspective. These new analyses allow us to evaluate previously untested questions, and examine the neural basis of flow from an entirely novel meta-theoretical and analytical paradigm. In what follows, we provide a brief overview of the procedure, before detailing the analytical approach.

#### 2.1.1 Participants

A total of *n* = 35 participants were recruited from The Ohio State University and the surrounding community (mean_age_ = 25.00, 45.71% female, 71.43% students, 48.57% Caucasian, and a mean self-reported video game ability of 4.11 out of 7.00). All participants were right-handed, had normal or corrected to normal vision, and did not demonstrate any contraindication to fMRI scanning. All study procedures were approved by The Ohio State University’s Institutional Review Board.

### 2.2 Stimulus and Experimental Manipulation

In the scanner, participants played a video game called *Asteroid Impact* following a previously validated experimental technique for inducing flow (Huskey et al., 2018). Specifically, flow is theorized to result from a balance between task difficulty and individual ability at the task (Csikszentmihalyi, 1990). Accordingly, we manipulated three experimental conditions: low-difficulty or boredom (difficulty < ability), high-difficulty or frustration (difficulty > ability), and balanced-difficulty or flow (difficulty = ability). Participants played all three conditions, and condition order was randomly assigned. In *Asteroid Impact,* players use a mouse cursor to control a spaceship with the goal of collecting targets on the screen while avoiding asteroids. Difficulty was manipulated by adjusting the speed of asteroids in one of three ways: (1) asteroid speed was slow and did not change (low-difficulty), (2) asteroid speed was consistently fast and did not change (high-difficulty), (3) an algorithm adjusted asteroid speed based on player performance (balanced-difficulty). A resting-state scan was also gathered after participants had played all three rounds of the game.

### 2.3 Dependent Measures

#### 2.3.1 Self-Report Measures: Flow and Enjoyment

The Autotelic Experience subscale of the Event Experience Scale was used to measure self-reported flow (Jackson & Eklund, 2004). Self-reported video game enjoyment was also measured (Bowman et al., 2013).

#### 2.3.2 Behavioral Measure: Reaction Time

Attentional engagement, a behavioral correlate of flow (for rationale and validation, see Huskey et al., 2018), was evaluated using a behavioral measure: reaction time (Lang et al., 2006). A total of 68 audiovisual (semi-opaque red circle; sine waveform, 440.0 Hz) probes were shown equally in each of the four corners of the game. Speeded responses were recorded using a previously validated technique (Calcagnotto et al., 2021).

### 2.4 fMRI Preprocessing

For complete functional magnetic resonance image (fMRI) acquisition and data cleaning details, we direct readers to (Huskey et al., 2022). In brief, all fMRI data were preprocessed using *fMRIPrep 20.0.5* (Esteban et al., 2019) and denoised using XCP engine (Ciric et al., 2018).

To optimize our data for the frequency domain analyses reported in this manuscript, we made two minor deviations to the data cleaning pipeline: (1) we used a slightly more permissive bandpass filter (0.01 - 0.08 *Hz* vs. 0.016 - 0.08 *Hz,* for further details, see below), and (2) we did not apply global signal regression (GSR; our previous analyses show trivial differences in quality control metrics when GSR is vs. is not applied to this dataset; for full details, see supplemental section 4 in Huskey et al., 2022).

### 2.5 Parcellation

We parcellated the pre-processed fMRI data by averaging time-courses across all voxels for each region defined in the anatomical parcellation AAL (Tzourio-Mazoyer et al., 2002) considering all cortical, subcortical, and cerebellar regions (*n* = 116). We chose the AAL parcellation as subcortical and cerebellar regions are relevant in flow studies (Gold & Ciorciari, 2021; Huskey et al., 2022).

### 2.6 Bandpass Filtering

To isolate low-frequency resting-state signal fluctuations, we bandpass filtered the parcellated fMRI time-series within 0.01-0.08 *Hz* with a discrete Fourier transform (DST) computed using a fast Fourier transform (FFT) algorithm in MATLAB 2021b. We applied Carson’s empirical rule (Carson, 1922; Pachaud et al., 2013) on the analytical signal which was calculated using the Hilbert transform of the real signal (Gabor, 1946), to confirm non-violation of the Bedrosian theorem for our band-passed signals.

### 2.7 Functional Connectivity Through Phase-Alignment

We estimated functional connectivity (FC) with the non-linear measure of phase-coincidence or phase-alignment which may be more suitable than linear measures, such as Pearson correlation, for analyzing complex brain dynamics. Specifically, non-linear methods provide insight into interdependence between brain regions at both short and large time and spatial scales allowing the analysis of complex non-linear interactions across space and time (Pereda et al., 2005; Quian Quiroga et al., 2002). From a practical perspective, unlike correlation or covariance measures, phase alignment can be estimated at the instantaneous level and does not require time-windowing. When averaged over a sufficiently long-time window, phase-alignment values provide a close approximation to Pearson correlation, varying within the same range of values (Cabral et al., 2017; Honari et al., 2021).

Following Cabral et al. (2017), we first calculated the analytical signal using the Hilbert transform of the real signal (Gabor, 1946). Then, the instantaneous phase-alignment between each pair of brain regions was estimated for each time-point as the cosine difference of the relative phase. Phase-alignment at a given timepoint ranges between −1 (regions in anti-phase alignment) and 1 (regions in phase alignment). A three-dimensional tensor of size *NxNxT* where *N* is the dimension of the parcellation, and *T* is the number of timepoints in the scan, was extracted for each participant for each condition.

### 2.8 LEiDA – Leading Eigenvector Dynamic Analysis

To reduce the dimensionality of the instantaneous phase-alignment (iPA) space for our dynamic analysis, we employed the Leading Eigenvector Dynamic Analysis (LEiDA; Cabral et al., 2017) method. The leading eigenvector *V_1_(t)* of each *iPA(t)* is the eigenvector with the largest magnitude eigenvalue and reflects the dominant FC (through phase-alignment) pattern at time *t*. *V_1_(t)* is a *Nx1* vector that captures the main orientation of the fMRI signal phases over all anatomical areas. Each element in *V_1_(t)* represents the projection of the fMRI phase in each region into the leading eigenvector. When all elements of *V_1_(t)* have the same sign, this means that all fMRI phases are orientated in the same direction as *V_1_(t),* indicating a global mode governing all fMRI signals. When the elements of *V_1_(t)* have both positive and negative signs, this means that the fMRI signals have different orientations behaving like opposite anti-nodes in a standing wave. This allows us to separate the brain regions into two ‘communities’ (or poles) according to their orientation or sign, where the magnitude of each element in *V_1_(t)* indicates the strength of belonging to that community (Newman, 2006). Complete details and graphical representations for this procedure are well documented (Figueroa et al., 2019; Lord et al., 2019; Vohryzek et al., 2020). The outer product of *V_1_(t)* reveals the FC matrix associated with the leading eigenvector at time t.

Community membership changes spontaneously or in response to a task. As these communities are based on relative phase, they may be viewed through the lens of Coordination Dynamics (Bressler & Kelso, 2001, 2016), and thus, patterns of community membership reflect different modes of coordination.

#### 2.8.1 Coordination Mode Extraction

To identify recurring spatiotemporal modes for phase-alignment or coordination patterns (modes, a term borrowed from physics, are different patterns of brain waves with the same frequency and a fixed phase relationship), we clustered the leading eigenvectors for each of the 4 time-series (flow, boredom, frustration, resting state) with k-means clustering with 300 replications and up to 400 iterations for 5 centroids. We choose 5 centroids as these spatiotemporal patterns have been shown to exhibit extremely high test-retest reliability in healthy subjects (Hancock, Cabral, et al., 2022). K-means clustering returns a set of K central vectors or centroids in the form of *Nx1* vectors *V_c_*. As *V_c_* is a mean derived variable, it may not occur in any individual subject data set. To obtain time courses related to the extracted coordination modes (*ψ_k_*) at each TR we assign the cluster number to which *V_c_(t)* is most similar using the cosine distance.

#### 2.8.2 Coordination Mode Representations as Connectograms

We visualized functional connectivity as connectograms by taking the functional connectivity matrices for each coordination mode and retaining regions that were in anti-phase alignment.

### 2.9 Neurosynth Functional Associations

Probabilistic measures of the association between brain coordinates and overlapping terms from the Cognitive Atlas (Poldrack et al., 2011) and the Neurosynth database (Yarkoni et al., 2011) were obtained as previously demonstrated in Hansen et al. (2021). The probabilistic measures were parcellated into 116 AAL regions and may be interpreted as a quantitative representation of how regional fluctuations in phase-alignment are related to psychological processes. The resulting functional association matrix represents the functional relatedness of 130 terms to 116 brain regions.

### 2.10 Complexity Dynamics of Global and Local Coordination

Signatures of complex phenomena extracted from neuroimaging data have been developed and estimated from the early 2000’s. However, even for one of the most popular signatures, metastability, agreement on what and how to estimate this signature is heavily debated (Hancock et al., 2025). Therefore, in the absence of a gold standard measure for complexity in the brain, we adopted a data driven approach and estimated several signatures that have been proposed in the literature to investigate which may be sensitive to behavioral phenomena, while also gaining clarity on the specific coordination modes associated with flow.

Within this framework, we distinguish between global and local coordination. Global coordination metrics collapse across regions to characterize macroscopic properties of whole-brain dynamics. Accordingly, these metrics describe how coordinated the brain is overall without reference to specific spatial patterns. By comparison, local coordination modes capture which regions coordinate together at a given moment, reflecting transient, recurring spatiotemporal patterns of phase alignment rather than static networks. An analogy would be considering brain activity like an orchestra where global coordination involves both the melody and the accompaniment, whereas local coordination involves just the melody.

Considering both global and local coordination is valuable because similar global dynamical regimes can arise from distinct underlying coordination patterns, and local modes provide insight into how whole-brain dynamics are instantiated at the regional level (for an extended discussion, see e.g., Bassett & Gazzaniga, 2011; Sporns, 2011). From a complex-systems perspective, these analyses operate at complementary levels of abstraction. This approach captures hierarchical organization in which macroscopic system-level dynamics emerge from, but are not reducible to, transient patterns of regional coordination (Anderson, 1972; Simon, 1962). Guided by this multilevel view of brain organization, we therefore quantify complexity at both the global and local levels using multiple established signatures drawn from the literature. Analytical aspects and a complete description of the global and local complexity signatures may be found in Supplementary Methods. We include here signatures that have been reported in the flow literature and those that yielded statistically significant results during our planned condition contrasts.

#### 2.10.1 Overview of Significant Signatures of Complexity for Global and Local Coordination

Synchronization measures the degree of synchronization between neural regions across the whole brain. It is calculated as the mean-field phase, which is more commonly known as the average of the Kuramoto Order Parameter (Shanahan, 2010). The synchronization measure is bound between [0, 1].

Metastability quantifies variability in synchronization, reflecting the switching between communities (i.e., regions in sync), and has been proposed as a key signature of metastability (Deco & Kringelbach, 2016). It is defined as either the variance or the standard deviation of Kuramoto Order Parameter, which is bound between [0, 0.272 (var), 0.552 (std)], and takes the mean-field phase into consideration (Shanahan, 2010).

Coalition Diversity measures the diversity of the communities that form through synchronization. It is calculated as the variance or the standard deviation of synchronization across all regions at each time point, averaged over the entire scan. Coalition diversity, also sometimes referred to as chimerality or a chimera state (Abrams & Strogatz, 2004), is bounded between [0, 0.272 (var), 0.552 (std)] (Shanahan, 2010).

Dynamical Alignment is a measure of the number of regions in anti-phase alignment, thereby indicating the extent of coordination dynamics. It is calculated through spectral decomposition and is measured as the average spectral radius or first eigenvalue of the complete phase alignment matrix over time. It is bound between [*0,N*] where *N* represents the number of regions in the parcellation (de Alteriis et al., 2024). If statistical relationships rather than phase relationships are used to derive the FC matrix, spectral decomposition will result in *N* eigenvectors and *N* eigenmodes. Hence, if the spectral radius is small, it indicates that the FC matrix has high dimensionality (i.e., only a small amount of variance in the FC matrix is explained by the first eigenvector). However, with phase alignment, there are just 2 eigenvectors and 2 eigenvalues. As a result, the spectral radius returns the number of regions in anti-phase alignment and its mean value across the duration of a scan provides an indication of the extent of regional coordination through anti-phase alignment (de Alteriis et al., 2024).

Entropy Rate (mean fMRI signal over all regions) is a newer complexity estimator for time series extending Lempel-Ziv (LZ) complexity. The entropy rate measures how many bits of innovation are introduced by each new data sample and is related to how hard it is to predict the next value of a sequence. This normalized LZ, *C*_LZ_, is a principled, data-efficient estimator of the diversity of the underlying neural process (Mediano et al., 2024). It is bound between [0,1].

Global Fluidity is the standard deviation of the functional connectivity dynamics (FCD). This signature reflects the fluidity of fluctuating internal coalitions (Petkoski et al., 2023). It is bound between [0,0.552]. FCD captures the spatiotemporal statistics of resting-state activity (E. C. A. Hansen et al., 2015).

### 2.11 Statistical Analysis

#### 2.11.1 Non-Parametric Statistical Tests

We investigated whether any of the complexity signatures differed statistically between task and rest, as well as among the flow, boredom, and frustration conditions. To compare these conditions, we used non-parametric repeated measures ANOVA (Friedman test) and 2-sided t-tests (Wilcoxon). This approach was chosen because most distributions were not normally distributed when assessed with a Shapiro test (see Supplementary tables ST2 and ST3).

#### 2.11.2 Linear Regression Modeling

We also sought to explore the explanatory power of complexity signatures in relation to subjects’ self-reported measures of flow and enjoyment, as well as the behavioral measure of reaction time. Before running our linear regression models, we removed correlations above 0.7 between the explanatory variables (complexity signatures) to avoid variance inflation due to multicollinearity (see Supplementary Figures SF1-SF3). To find the best-fitting models for explaining flow, enjoyment, and reaction time, we employed stepwise regression. That is, models were built with 1 to *n* non-correlated explanatory variables (complexity signatures). The average prediction error (RMSE) of the *n* best models was then estimated with 10-fold cross-validation with 3 repetitions (Kuhn & Johnson, 2013). The model that minimized the RMSE was selected. Linear modeling was performed with the Caret package (Kuhn, 2008) implemented in R.

#### 2.11.3 Familywise Error Rate Correction

To control for potentially inflated familywise error rates (FWER), we applied false discovery rate corrections (FDR; Benjamini & Hochberg, 1995; Benjamini & Yekutieli, 2001) to our statistical tests and models. We report unadjusted p-values alongside the corresponding FDR-adjusted significance thresholds (p-critical), with statistical significance determined when the unadjusted p-value fell below its associated FDR critical value. Importantly, we applied different thresholds for different analyses to reflect our distinct inferential goals. Analyses comparing task and condition effects on complexity signatures were designed to test a priori contrasts across a large number (90 global comparisons; 105 local comparisons) of predefined coordination metrics. These analyses therefore employed a more conservative FDR threshold (*q* = 0.05) to limit false discoveries in the context of planned comparisons.

In contrast, our linear regression analyses were explicitly exploratory. These models simply test if coordination metrics (9 global models; 45 local models) explain variance in subjective experience and behavior. Given our data-driven objective, and the additional safeguards provided by cross-validation and multicollinearity censoring, we applied a more permissive FDR threshold (*q* = 0.10), which is consistent with common practice in exploratory modeling contexts (for extended discussion, see Goeman & Solari, 2011).

This approach reflects the broader distinction between confirmatory inference, which prioritizes stricter error control, and exploratory inference, which prioritizes hypothesis generation while maintaining transparent and principled control of false discoveries (Goeman & Solari, 2011).

### 2.12 Software tools

Parcellation, LEiDA, and all complexity signature calculations were implemented in MATLAB (MATLAB, 2021). Neurosynth functional associations were derived in Python 3.8.5. All other statistical analyses were performed in RStudio Team version 2023.09.1 Build 494.

### 2.13 Open Science Practices, Supplemental Information

This manuscript conducts new analyses on data previously published by Huskey and colleagues (2022), who made the raw functional magnetic resonance imaging data available on OpenNeuro (https://openneuro.org/datasets/ds003358/versions/1.0.0) and the raw self-report and behavioral data are posted to the Open Science Framework (OSF; https://osf.io/bxvhr/). The experimental task, *Asteroid Impact*, is available on GitHub https://github.com/asteroidimpact. New code was written to analyze these data, which is available on GitHub (https://github.com/franhancock/COMPLEXITY_GLOBAL_AND_LOCAL). Supplemental information referenced in this manuscript are posted to OSF (https://osf.io/f73qm/).

## 3. Results

### 3.1 Global Complexity Signatures

We calculated 15 global signatures of complexity (see Supplementary Methods), each characterizing different aspects of whole-brain dynamics, for the four studied conditions: flow, boredom, frustration, resting state. A non-parametric Friedman test (for repeated measures) was applied to assess differences between conditions in all 15 complexity signatures. Following FDR correction, 27 of the 90 pairwise comparisons remained statistically significant (Supplementary Table ST2, Supplementary Figure SF7). Discarding differences that were nonspecific to our phenomenological manipulations and, therefore, likely understood as general effects of gameplay (i.e. where differences were found in multiple task-based conditions when compared to rest), seven significant results in four complexity signatures remained (Figure 1).

**Figure 1.**
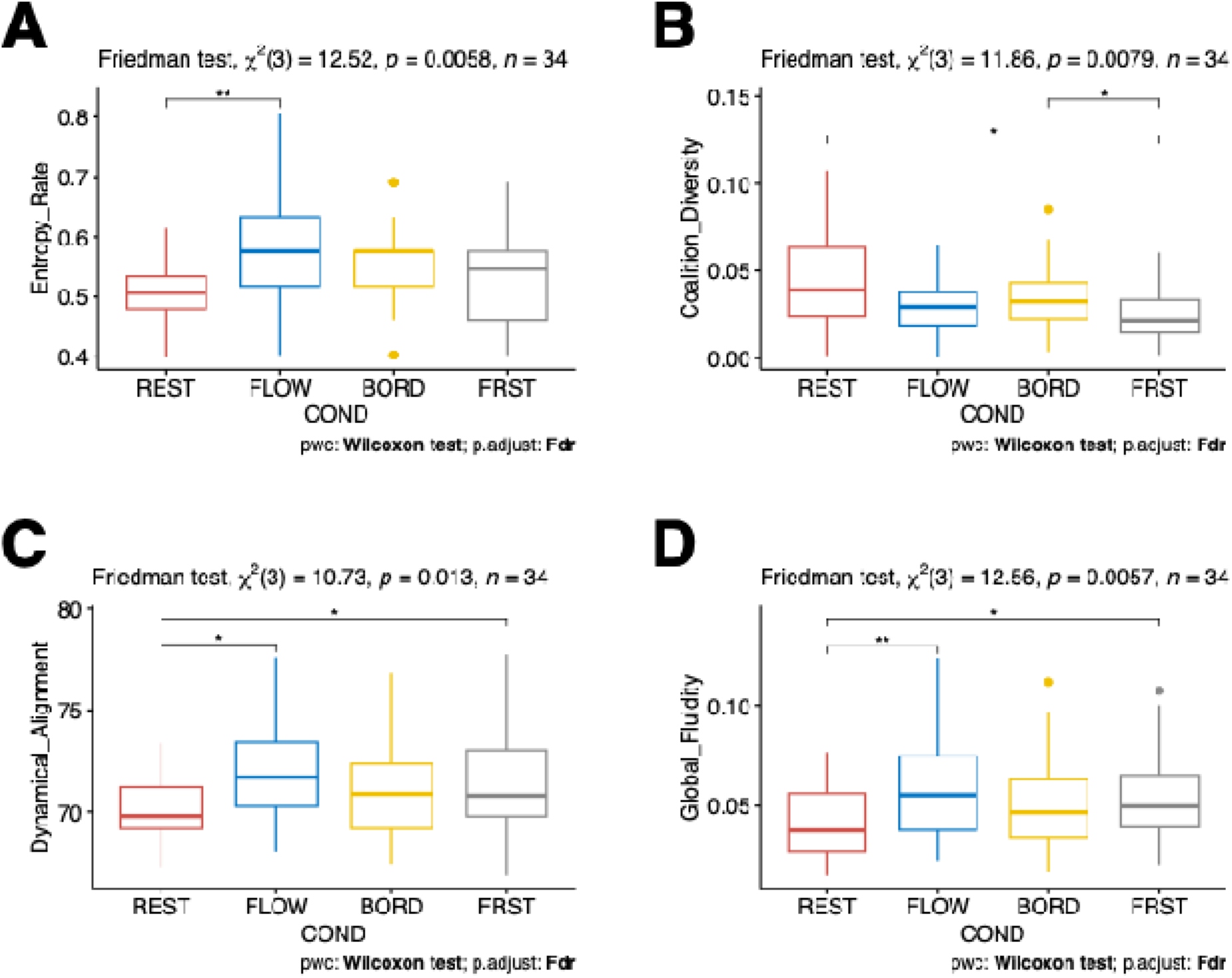
Non-gameplay related significant difference in Global Complexity Signatures. A) Entropy Rate. B) Coalition diversity. C) Dynamical Alignment. D) Global Fluidity. \**p*< 0.05; \*\**p*<0.01; \*\*\**p*<0.001.

The first signature was entropy rate (Figure 1A; *χ*^2^(3)=12.52, *p_j_*=0.0058). A post-hoc pairwise comparison indicated that this difference specifically pertained to higher entropy rate values in the flow condition compared to rest (*W*=83, *p*<0.001, *p_crit_*=0.01, *effect size*=0.514).

The second was Coalition Diversity (Figure 1B; χ^2^(3)=11.86, *p*=0.0079). Post-hoc pairwise comparison revealed that this difference related to high Coalition Diversity values in the boredom condition compared to the frustration condition (*W*=440, *p*=0.014, *p_crit_*=0.015, *effect size*=0.306) and in the rest condition compared to the frustration condition (*W*=470, *p*=0.002, *p_crit_*=0.012, *effect size*=0.373).

The third signature was Dynamical Alignment (Figure 1C; χ^2^(3)=10.73, *p*=0.013). Post-hoc pairwise comparison showed that this difference was related to higher values in the flow condition compared to rest (*W*=142, *p*=0.007, *p_crit_*=0.013, *effect size*=0.358), and in the frustration condition compared to rest (*W*=150, *p*=0.011, *p_crit_*=0.014, *effect size*=0.268).

Finally, the fourth signature was Global Fluidity (Figure 1D, χ^2^(3)=12.57, *p*=0.006). Post-hoc pairwise comparison revealed that this difference was related to higher values in the flow condition compared to rest (*W*=117, *p*=0.001, *p_crit_*=0.011, *effect size*=0.358), and in the frustration condition compared to rest (*W*=135, *p*=0.005, *p_crit_*=0.013, *effect size*=0.297).

### 3.2 Local Coordination Complexity Signatures

Evaluating complexity signatures in local coordination modes (see Methods) enables a more precise and comprehensive interrogation of brain dynamics during boredom, flow, and frustration, complementing our global analyses. The coordination modes are examined descriptively and evaluated inferentially in Supplementary Methods. Here we present the associated figures to facilitate an applied understanding of the analytical results (Figure 2) and the cognitive, affective, and motor processes that characterize each local coordination mode (Figure 3). Together, these figures render abstract coordination modes concrete by linking statistical structure to functional brain organization.

**Figure 2.**
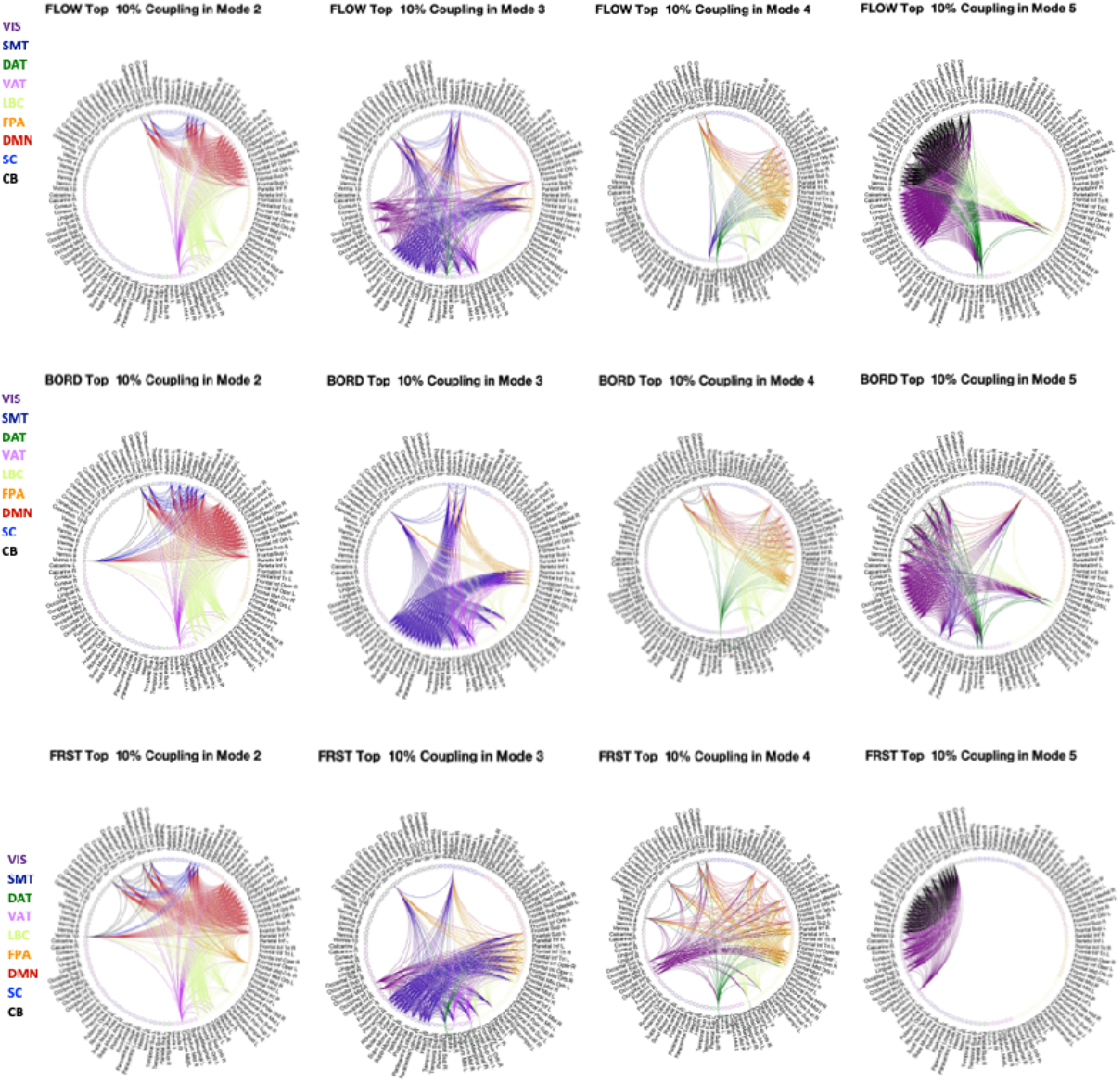
Modes of coordination based on phase alignment. VIS, visual network; SMT, somatomotor network; DAT, dorsal attention network; VAT, ventral attention network; LBC, limbic network; FPA, frontoparietal; DMN, default mode network; SC, subcortical; CB, cerebellum. Descriptions may be found in the Supplementary Methods.

**Figure 3.**
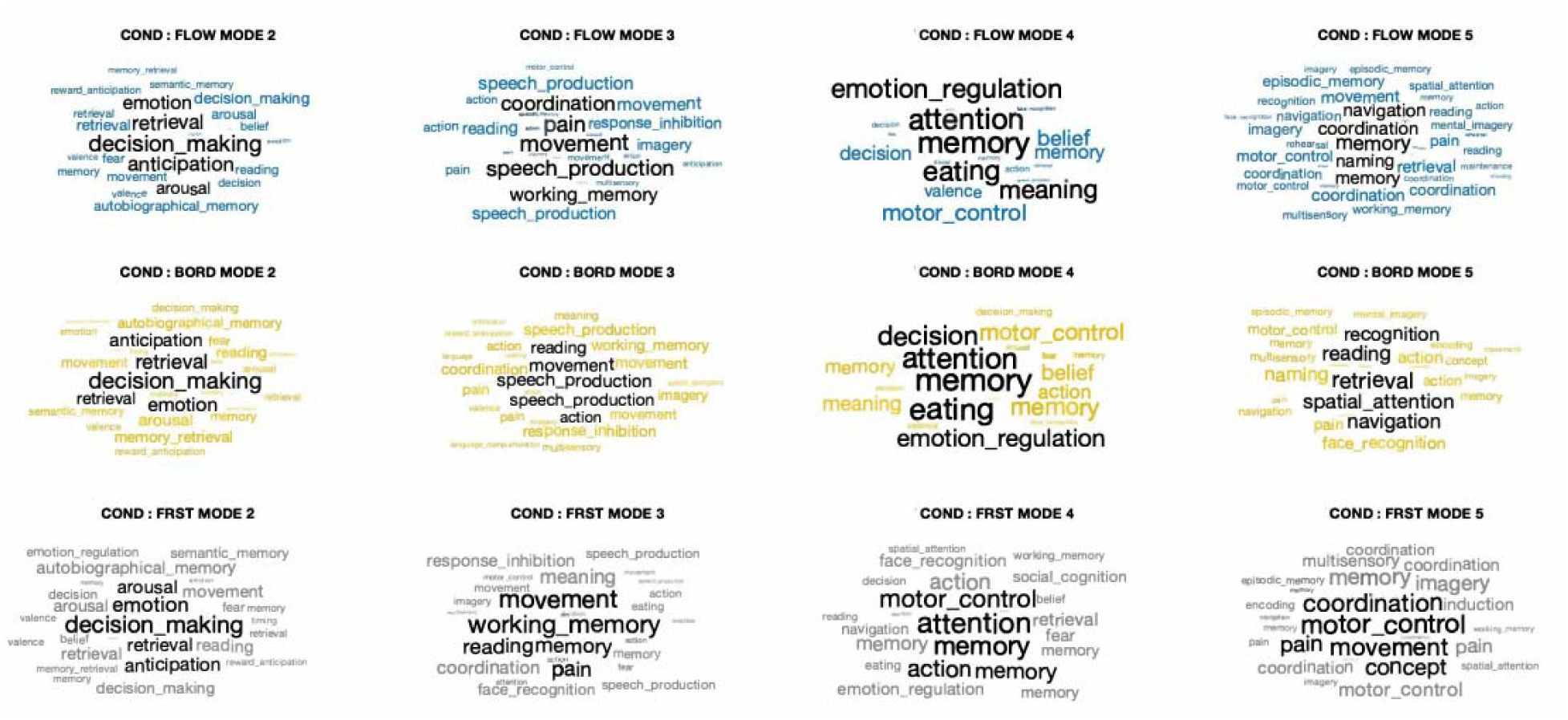
Neurosynth meta-analysis of the coordination modes. The psychological concepts associated with the brain regions in each coordination mode are presented here in a word-cloud format. Word size is proportional to the region’s contribution to the phase alignment of the mode.

We estimated the coordination mode level signatures of duration, occurrence, synchronization, metastability, dynamical complexity, dynamical alignment, and dynamical agility across the three task conditions. Following FDR correction (q = 0.05) for multiple comparisons, we found that 1 of 105 comparisons remained statistically significant. Specifically, Dynamical Alignment (i.e., increased order) in coordination mode 4 was higher in the boredom condition relative to the frustration condition (Figure 4; *W* = 879, *p<0.001, p_crit_=*0.0004, *effect size*=0.448). Although Dynamical Alignment in coordination mode 4 was also significantly higher in the flow condition compared to the frustration condition with FDR applied to just this signature, following FDR for all signatures, in all modes, across all conditions, the difference did not remain statistically significant (*W*=814, *p*=0.004, *p_crit_*=0.001, *effect size*=0.351). See Supplementary Table ST3 and Supplementary Figure ST8 for all results and calculation of FDR.

**Figure 4.**
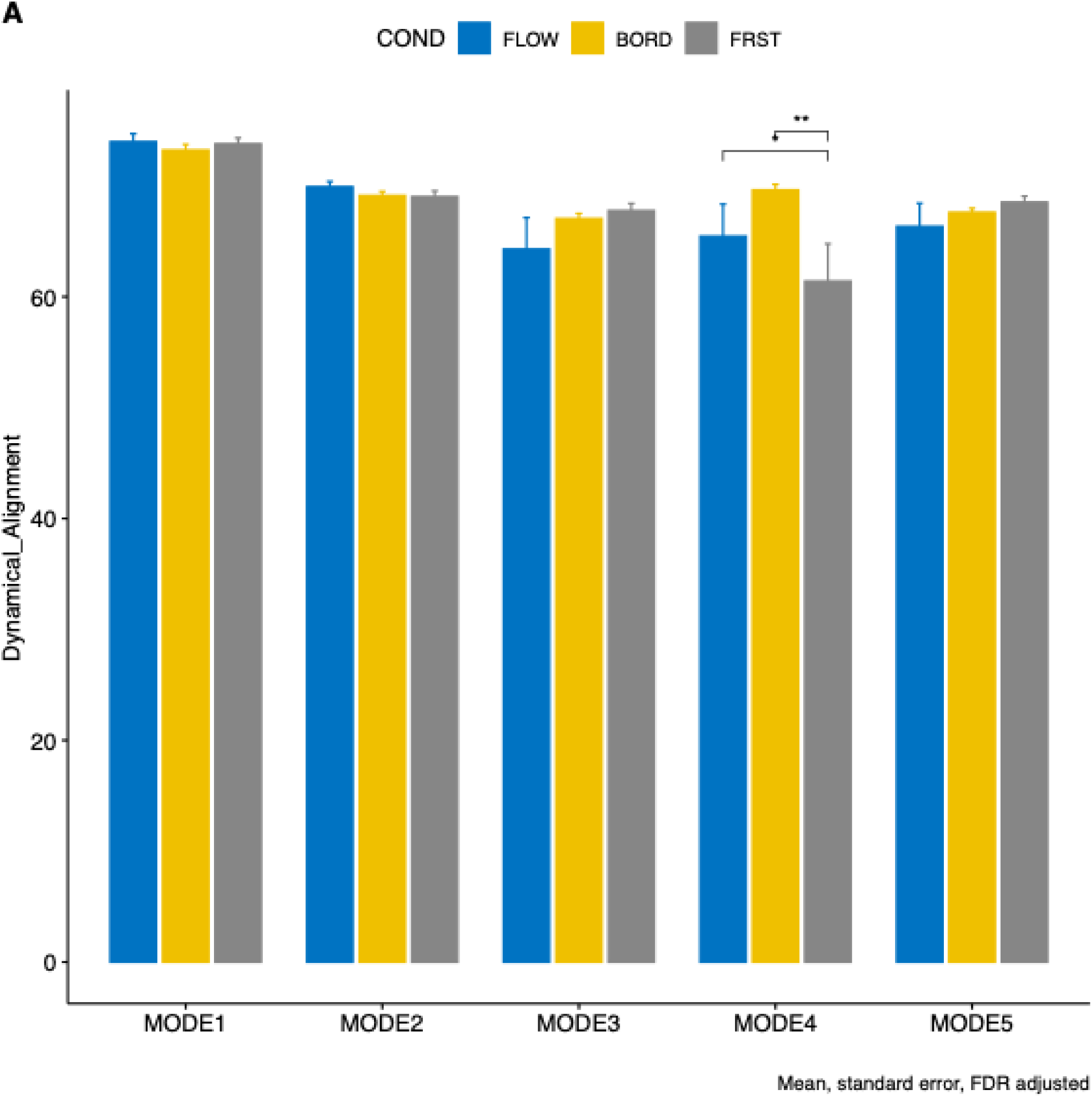
Differences in local complexity signatures, dynamical alignment. Note that the difference between FLOW and FRST did not survive FDR for pairwise comparisons across all 105 comparisons. FLOW, flow condition; BORD, boredom condition; FRST, frustration condition. \**p*< 0.05; \*\**p*<0.01; \*\*\**p*<0.001.

### 3.3 Examining Brain-Self-Report and Brain-Behavioral Relationships

Explanatory data analysis was used to capture relationships between complexity signatures and the self-report (i.e., flow, enjoyment) and behavioral (i.e., reaction time) scores in the flow, boredom, and frustration conditions. Our rationale was aimed at identifying differences between the conditions that could potentially disambiguate brain dynamics during flow, boredom, and frustration. Following FDR correction, we found statistical significance in 4 of the 54 models.

Results are summarized in Table 1, whereas comprehensive model results and FDR results may be found in the Supplementary tables ST4-12. Statistically significant models that produced the highest coefficient of variation for each score type in each condition are reported below.

**Table 1.** Significant contributions of complexity signatures in linear modes.

|  |  |  |  |  |  |  |
| --- | --- | --- | --- | --- | --- | --- |
| Experimental Condition: Flow |  |  |  |  |  |  |
| Global | Outcome Variable | $R^2$ | $R^{2\text{ Adj}}$ | p-value | p-critical | Explanatory Terms |
|  | Flow | 0.19 | 0.18 | 0.0106 | 0.0111 | Entropy Rate |
| Experimental Condition: Boredom |  |  |  |  |  |  |
| Local | Outcome Variable | $R^2$ | $R^{2\text{ Adj}}$ | p-value | p-critical | Explanatory Terms |
| Mode 3 |  |  |  |  |  |  |
|  | Flow | 0.44 | 0.34 | 0.004 | 0.007 | Synchronization, Metastability, Dynamical complexity, Duration, Dynamical agility |
|  | Enjoyment | 0.38 | 0.32 | 0.002 | 0.004 | Synchronization, Metastability, Dynamical agility |
| Experimental Condition: Frustration |  |  |  |  |  |  |
| Local | Outcome Variable | $R^2$ | $R^{2\text{ Adj}}$ | p-value | p-critical | Explanatory Terms |
| Mode 2 |  |  |  |  |  |  |
|  | Enjoyment | 0.44 | 0.38 | 0.001 | 0.002 | Synchronization, Metastability, Dynamical agility |

#### 3.3.1 Best Explanatory Models in the Flow Condition

##### Flow

The best model explained 19% (18% adj, *p*=0.0106, *p_crit_* =0.011) of the variance in flow scores. Entropy rate made a statistically significant contribution to the model, such that for any unit decrease in entropy rate, the model explained higher flow scores.

##### Enjoyment

No model explained the variance in enjoyment scores.

##### Reaction time

No model explained the variance in the reaction time scores.

#### 3.3.2 Best Explanatory Models in the Boredom Condition

##### Flow

The best model explained 44% (34% adj., *p*=0.004, *p_crit_*=0.007) of the variance in flow scores. Synchronization and metastability in coordination Mode 3 made statistically significant contributions: while holding dynamical complexity, duration, and dynamical agility constant, higher flow scores were associated with decreased synchronization and increased metastability.

##### Enjoyment

The best model explained 38% (32% adj., *p*=0.002, *p_crit_*=0.004) of the variance in enjoyment scores. Synchronization and metastability in Mode 3 made statistically significant contributions to the model. When holding dynamical agility constant, the model explained higher scores for enjoyment for any unit decrease in synchronization or any unit increase in metastability.

##### Reaction time

No model explained the variance in the reaction time scores.

#### 3.3.3 Best Explanatory Models in the Frustration Condition

##### Flow

No model explained the variance in the flow scores.

##### Enjoyment

The best model explained 44% (38% adj., *p*=0.001, *p_crit_*=0.002) of the variance in enjoyment scores. Synchronization and metastability in Mode 2 made statistically significant contributions to the model. When holding dynamical agility constant, the model explained higher scores for enjoyment for any unit decrease in synchronization or any unit increase in metastability.

##### Reaction time

No model explained the variance in the reaction time scores.

### 3.4 Global entropy rate

Although global entropy rate was not significantly different between flow and the boredom or frustration conditions, it nevertheless explained a moderate proportion of the variance in the flow scores in the flow condition. We therefore investigated if the relationships between the flow scores and global entropy differed across conditions. To do this, we tested if there were significant differences in either the intercepts or the slopes of a simple regression model using centered flow scale data to ensure that interpretability of the results. We found that the slopes did not significantly differ (the interaction terms are not statistically significant), but the intercept of the flow condition was significantly higher than the boredom condition as indicated in the regression output in Table 2 and visible in the regression plot, Figure 5. Therefore, at the mean of the flow scores, global entropy rate appears to be statistically higher in the flow condition than in either the boredom or frustration condition.

**Figure 5.**
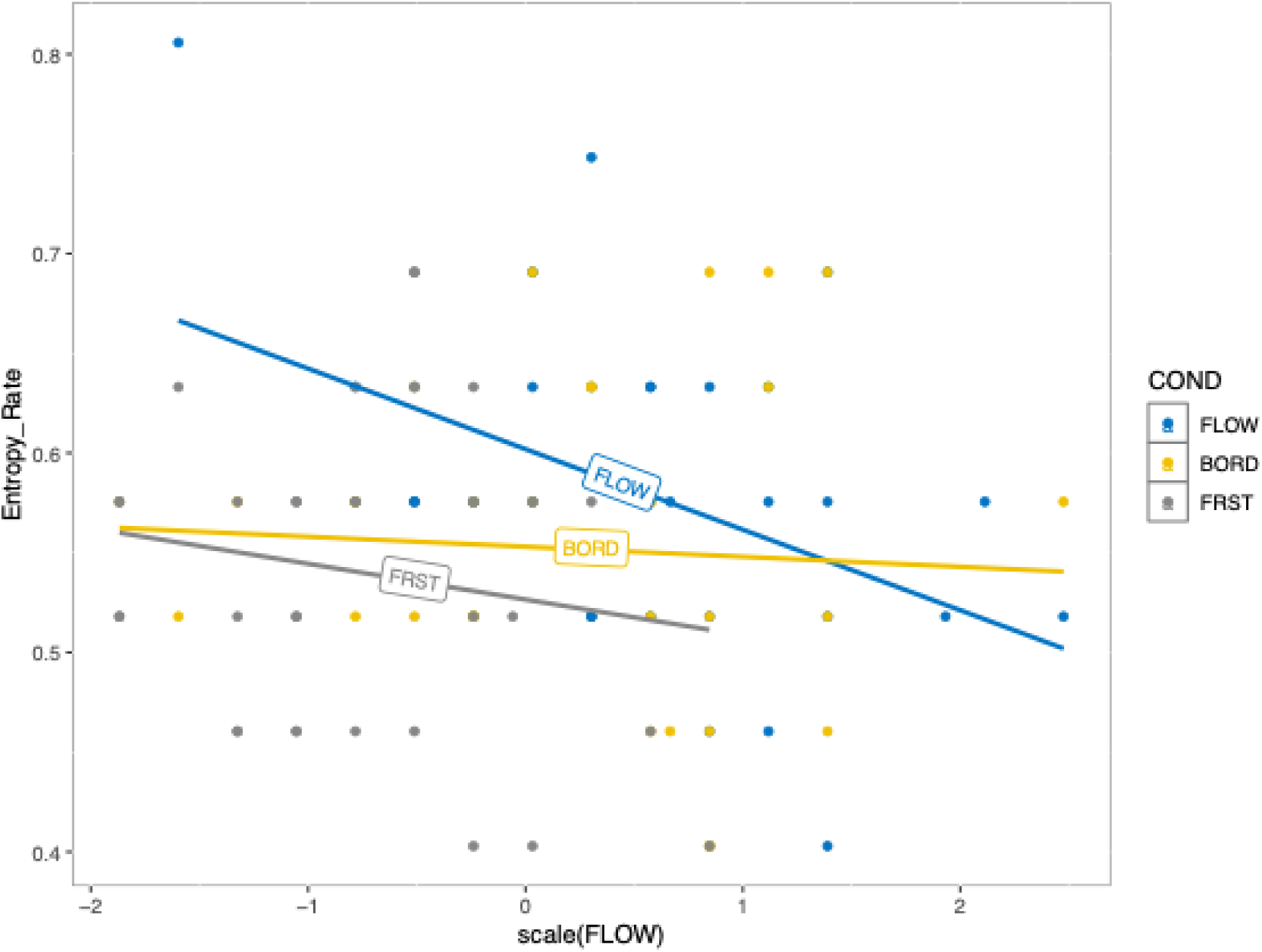
Regression of centered flow scores against entropy rate across the 3 conditions. In this analysis, the intercept represents the mean value of global entropy rate at the mean value of the flow scores.

**Table 2.** Output from the linear regression of flow scores against entropy rate in all 3 conditions.

| Residuals |  |  |  |  |  |
| --- | --- | --- | --- | --- | --- |
|  | Min | 1Q | Median | 3Q | Max |
|  | -0.14585 | -0.047004 | -0.00178 | 0.048404 | 0.158564 |
| Coefficients |  |  |  |  |  |
|  | Estimate | Std. Error | t value | Pr(>\|t\|) |  |
| (Intercept) | 0.72127 | 0.05686 | 12.685 | <2e-16*** |  |
| FLOW | -0.04386 | 0.01723 | -2.546 | 0.0125* |  |
| CONDBORD | -0.15337 | 0.07419 | -2.067 | 0.0414* |  |
| CONDFRST | -0.1418 | 0.07267 | -1.951 | 0.0539 |  |
| FLOW:CONDBORD | 0.03839 | 0.02325 | 1.651 | 0.1019 |  |
| FLOW:CONDFRST | 0.0244 | 0.02741 | 0.89 | 0.3755 |  |
| *$p<0.05$; **$p<0.01$; ***$p<0.001$ | | | | | |
| Residual standard error: 0.07862 on 96 degrees of freedom |  |  |  |  |  |
| Multiple R<sup>2</sup>: 0.1131, Adjusted R<sup>2</sup>: 0.06688 |  |  |  |  |  |
| $F$-statistic: 2.448 on 5 and 96 DF, $p$-value: 0.03919 | | | | | |

### 3.5 Examining Hypo-Frontality

The hypo-frontality (Dietrich, 2004) hypothesis has received mixed support over the years (Harris et al., 2017; Kotler et al., 2022) and has not been evaluated from a complexity science perspective. Therefore, we examined our data for evidence in support of the idea. Operationalizing hypo-frontality at the regional level, however, requires a principled mapping of the dorsolateral prefrontal cortex (DLPFC) within a standardized anatomical framework. The regions of the DLPFC are, unfortunately, not clearly defined in MNI coordinates (Pàmies-Vilà et al., 2023). We used the study of Limb & Braun (2008) who documented MNI coordinates for the bilateral medial DLPFC, lateral DLPFC, and superior DLPFC, and observed deactivations in each during flow, to map the DLPFC to AAL116 regions. We identified bilateral Frontal Mid (middle frontal gyrus – L – memory and retrieval, R – memory and cognitive control) and Frontal Inf Tri Right (inferior frontal gyrus, triangular part – empathy and meaning) as the AAL116 atlas equivalents (Table 3).

**Table 3.**
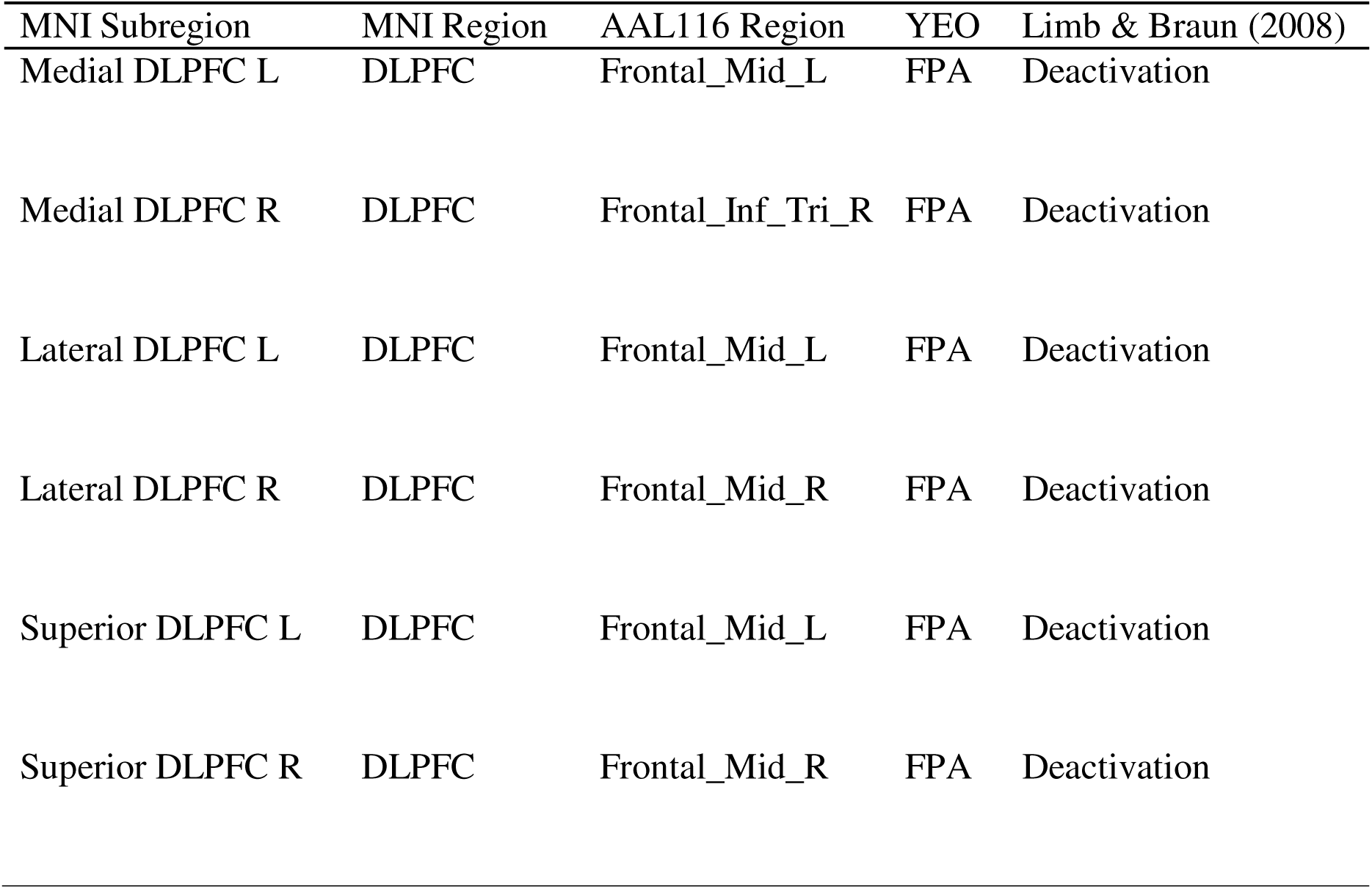
Mapping of AAL116 ROIs to MNI coordinates.

| MNI Subregion | MNI Region | AAL116 Region | YEO | Limb & Braun (2008) |
| --- | --- | --- | --- | --- |
| Medial DLPFC L | DLPFC | Frontal_Mid_L | FPA | Deactivation |
| Medial DLPFC R | DLPFC | Frontal_Inf_Tri_R | FPA | Deactivation |
| Lateral DLPFC L | DLPFC | Frontal_Mid_L | FPA | Deactivation |
| Lateral DLPFC R | DLPFC | Frontal_Mid_R | FPA | Deactivation |
| Superior DLPFC L | DLPFC | Frontal_Mid_L | FPA | Deactivation |
| Superior DLPFC R | DLPFC | Frontal_Mid_R | FPA | Deactivation |

Inspecting the contribution of these regions to the coordination modes, we found that all 3 regions contributed to coordination Mode 4 (attention) but were absent in coordination Mode 3 (movement) in the flow condition. In contrast Frontal Mid L (memory and retrieval) did not contribute to any coordination mode in the boredom condition, Frontal Mid R (memory and cognitive control) contributed to coordination Mode 4 (attention), and Frontal Inf Tri R (empathy and meaning) contributed to both coordination Mode 3 (movement) and 4 (attention). In the frustration condition, only Frontal Mid L (memory and retrieval) contributed to coordination Mode 4 (attention), whilst the other two regions (memory and cognitive control, empathy and meaning) contributed to coordination Mode 3 (movement) (see Figure 6).

**Figure 6.**
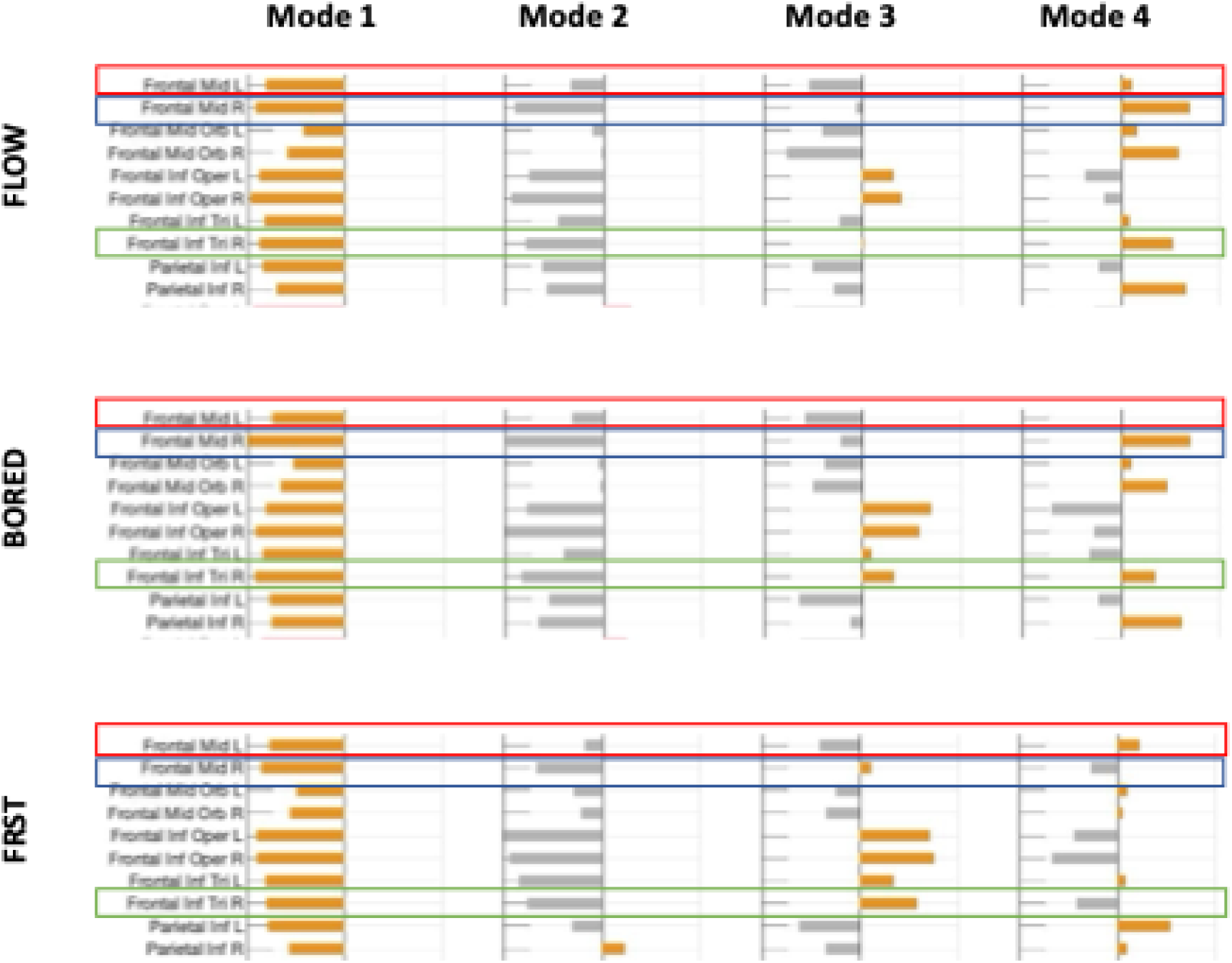
Partial hypo-frontality in the flow, boredom, and frustration conditions. Details of the regions contributing to coordination dynamics for the Frontal Parietal Network in all three conditions. Dorsolateral prefrontal cortex (DLPFC) regions are indicated with red, blue and green bars.

It appears that in the flow condition, hypo-frontality of subregions of the DLPFC exists during periods when coordination Mode 3 is dominant (i.e., during periods associated with movement). The DLPFC, however, contributes to coordination dynamics in the flow condition when coordination Mode 4 is dominant (i.e., during periods associated with attention).

## 4. Discussion

To date, the exploration of the neural dynamics of flow has been dominated by reductive (i.e., locationist, connectionist) theories. A complexity science approach, however, captures the multiple and interacting cognitive processes that underlie the flow experience, and offers a perspective the locationist and connectionist paradigms cannot provide. This study extends flow research in this direction by applying a battery of complex systems analyses to characterize three phenomenological experiences: flow, boredom and frustration. More specifically, we analyzed the differences between complexity signatures of coordination dynamics at the global and local (mode) level across four conditions and explored the explanatory power of 15 global and 7 local signatures for self-reported and behavioral scores. Our results add nuance to, and in some cases challenge, existing neural models of flow.

### 4.1 The Neural Complexity Signatures of Flow, Boredom, and Frustration

Our results revealed, at the global coordination level, that entropy rate during the flow condition was higher than at rest, was inversely proportional to subjective flow scores in the flow condition, and the intercept for the flow condition was statistically significantly higher than for the boredom condition. Additionally, we found similarities between the flow and frustration conditions. Specifically compared to rest, both demonstrated a larger number of regions involved in coordination dynamics (indexed by dynamical alignment), constantly switching (indexed by global fluidity) between a set of communities. The communities were more diverse (indexed by coalition diversity) in both the rest and boredom conditions relative to the frustration condition, but there was no difference in diversity between the flow and frustration condition.

Intriguingly, at the coordination mode level, dynamical alignment was higher in the boredom condition than the frustration condition in coordination mode 4. This means many regions were involved in these coordination dynamics. Together, the positive association between metastability and flow scores in the boredom condition together with the diverse (indexed by coalition diversity) and large set of regions involved in the partial synchronization (indexed by dynamical alignment), appear to indicate that flow scores in the boredom condition benefit from constantly changing coordination across a large set of diverse brain regions. This may be the reason that synchronization and metastability in the boredom condition explained flow scores at the coordination mode level (coalition diversity is based on synchronization over space), whereas this was not the case with the flow or the frustration conditions.

We note that increased enjoyment scores for the boredom and frustration conditions were explained by decreasing synchronization and increasing metastability, but in different coordination modes. In the case of boredom, the enjoyment association was found in the coordination mode associated with movement and working memory, whereas in the case of frustration, the association was found in the coordination mode associated with decision-making, anticipation, and emotion. A possible explanation could be that in the boredom condition, the participants enjoyed switching between recalling memorized movements and their execution, whereas in the frustration condition, enjoyment was increased when participants did not get ‘stuck’ (i.e. synchronized) in either decision-making, anticipation, or emotion. We now discuss these results in greater detail, below.

#### 4.1.1 Entropy Rate

The finding that entropy rate was significantly higher in the flow condition than in rest is in alignment with previous literature. Prior research notes that flow is an altered state of consciousness that may share some neural dynamics commonalities with other altered states such as the psychedelic state (Kotler et al., 2022). Increases in entropy rate have been associated with a variety of altered states of consciousness (Mediano et al., 2024; Vivot et al., 2020) including, during the acute psychedelic experience (Schartner et al., 2017; Timmermann et al., 2019), meditation (Vivot et al., 2020), and music improvisation (Dolan et al., 2018). Entropy rate has also been shown to increase during task performance, and, notably, elevated entropy rate, as reported by Dolan et al. (2018) and Rosen et al. (2024), has been linked to subjective experiences of “creative flow” during music improvisation. This interpretation aligns with evidence that higher entropy states are perceived as subjectively *richer* experiences (Bonnelle et al., 2024; Carhart-Harris, 2018).

However, the finding that flow was inversely proportional to entropy rate was perplexing. Inspecting Figure 5, we see that entropy rate in the flow condition falls to the rate at rest (*m=*0.507*, sd=*0.045, see Supplementary Table ST2, Tab ‘descriptives’) when flow scores increase to 4. An explanation for this unexpected behavior may lie in the experimental setup. According to Mediano et al. (2024), a form of “complexity matching” (Carpentier et al., 2020) may occur where neural dynamics are entrained by an external stimulus, in our case the video game, which obscures the relationship between neurodynamics and subjective experiences. They found that increased within-brain correlations driven by external simulation could obscure potential correlations between entropy rate and individual scores. Additionally, they found that subjects (under LSD) watching a video had the highest absolute entropy rate but did not give maximal subjective ratings. We may have observed a similar result in our dataset.

Alternatively, given the nature of the presently used tasks, our results suggest a potential distinction between flow states occurring during highly creative tasks where higher entropy rate is conducive to flow, and those occurring during less-creative tasks (as in the present case) where higher entropy rate appears detrimental to flow (for a discussion, see e.g., van der Linden et al., 2021). We note, however, that this interpretation is highly speculative, and deserving of examination in future research.

#### 4.1.2 Synchronization and Metastability

Synchronization between brain networks (i.e., attention and reward, fronto-parietal and reward) has been proposed to underlie the flow experience (Huskey et al., 2018; Weber et al., 2009), and more recent theorizing has speculated that metastable brain dynamics may underlie flow (Kotler et al., 2022). Network neuroscience results supporting these hypotheses are mixed, and appear dependent on, among other things, analytical approach. When examined in the time domain, averaged across task, and linearly modeled using psychophysiological interaction analysis (PPI; Friston et al., 1997), it appears that flow is associated with functional connections between the nucleus accumbens (ventral striatum) and dorsolateral prefrontal cortex (Huskey et al., 2018). However, more recent work using instantaneous phase analyses shows that both synchronous and metastable connections between fronto-parietal and reward networks are lowest during flow, particularly when compared to frustration (Huskey et al., 2022).^3^

In this analysis, we find more complicated results linking connectivity between subcortical reward structures and attentional/fronto-parietal structures. In coordination Mode 2, we observe coupling between the bilateral caudate nucleus (dorsal striatum) and left cingulum mid (ventral attention), although we note that this coupling is observed across all three conditions, therefore it does not appear unique to flow (see Supplementary figures SF4-SF6). Additionally, in coordination Mode 2, the bilateral caudate (dorsal striatum) is broadly coupled with the default mode network, limbic system, and ventral attention network for flow, boredom, and frustration. Similarly, in coordination Mode 3, the right pallidum (basal ganglia) shows widespread coupling with ventral attention, visual, somatomotor, and frontal-parietal structures for both flow and boredom. In coordination Mode 3, we also see that the right putamen (dorsal striatum) shows widespread connectivity with dorsal and ventral attention, visual, somatomotor, and frontal-parietal structures for flow, boredom, and frustration. These findings demonstrate that connectivity between fronto-parietal and reward, or attention and reward networks is associated with, but not unique to, flow. This undermines a core premise of synchronization theory (Weber et al., 2009).

Similarly, when examining brain-phenomenology (i.e., self-report) or brain-behavior (i.e., reaction time) relationships, we do not see evidence that synchrony or metastability explains variance in phenomenology or behavior during the flow condition. Instead, and surprisingly, our linear regression models show that coordination Mode 3 synchronization is detrimental to both flow and enjoyment during the boredom condition. By comparison, increases in coordination Mode 3 metastability are associated with increased flow and enjoyment. For the frustration condition, we find that, for coordination Mode 2, decreases in synchronization are associated with increased enjoyment, whereas increases in metastability contribute positively to enjoyment.

It is important to contextualize these relationships between phenomenology across conditions. Self-reported flow and enjoyment are highest in the flow condition, followed by the boredom and then frustration conditions (see Table 2 in Huskey et al., 2022). The implication is that synchrony and metastability only explain variance in phenomenology during conditions that do not elicit high levels of flow; a finding that undermines theories that strongly implicate synchrony (Weber et al., 2009) or metastability (Kotler et al., 2022) in flow.

Nevertheless, the complexity signature patterns observed in the boredom and frustration conditions raise intriguing questions. Both synchronization and metastability explain flow and enjoyment in these conditions, but the coordination Modes engaged were distinct. In boredom, both synchronous and metastable signatures were associated with flow and enjoyment specifically in coordination Mode 3—a pattern linked to movement, working memory, and response inhibition. This may reflect the fact that, although participants reported boredom, they remained engaged in the task with a high degree of self-control, enabling seamless cognitive control. Flow has been linked to reward-modulated cognitive control (Huskey et al., 2018), and the boredom condition may have offered just enough demand to generate a subjectively enjoyable experience, even if that experience was less enjoyable than the flow condition.

Enjoyment in the frustration condition, alternatively, was characterized by synchronization and metastability in coordination Mode 2, a mode associated with emotion, decision making, and anticipation. As mentioned above, the enjoyment relationship with synchronization (negative) and metastability (positive) suggests that enjoyment increases when subjects do not dwell too long (synchrony) on emotions, decision-making or anticipation, but rather continuously switch (metastability) between these and other psychological thoughts when coordination mode 2 is dominant.

Overall, this pattern of results suggests that synchrony and metastability play important roles in the phenomenological experiences of boredom and frustration. This may be also the case for flow, but the effect may be overshadowed by entropy rate. Future research may more carefully interrogate the positive relationship between metastability and phenomenological markers during conditions of boredom or frustration. Notably, these phenomenological experiences are often experienced as aversive (K. A. Mathiak et al., 2013; Scheepers & Keller, 2022; Westgate, 2020). It could be that metastable brain dynamics facilitate flexibility in neural coupling (Tognoli & Kelso, 2014) preparing the brain to transition out of (rather than being locked into) these aversive experiences.

### 4.2 The Utility of a Complexity Science Approach

Within the framework of complexity science, we did not find any global or local complexity signature that distinguished flow from other phenomenological experiences following strict FDR corrections. We acknowledge that investigating so many metrics could appear like data-dredging, but one objective of the analysis was to understand the similarities and differences between the multitude of metrics in the literature for analyzing brain dynamics. Even with the correlation matrices for the complexity signatures (Supplementary Figures ST1:ST3), we were surprised to see that these differed across conditions which adds to the complexity of unravelling brain dynamics with a set of orthogonal metrics.

However, when we moved away from simple analysis of means to linear regression models of brain-phenomenology, we were able to link complexity signatures with self-reported flow. This point is particularly worth emphasizing as few papers explicitly link brain response to phenomenological experience (a longstanding limitation in this research area). In doing so, we gain new insights into the brain-network dynamics associated with flow. More broadly, these results support not only the dynamic nature of brain functioning, but also reject both locationist and simple connectionist models of brain function (for an extended argument, see Noble et al., 2024) by recognizing that individual brain regions repeatedly align to form a set of communities (coordination modes) rather than exclusively belonging to a single functional network. This means that the contribution of each brain region within a coordination mode changes over time, thereby enabling endless configurations both spontaneously (Hancock, Cabral, et al., 2022; Hancock et al., 2023) and for the first time now, in response to a task.

### 4.3 Implications for Theories of Flow

Our findings challenge and extend existing theories of flow by demonstrating that no single, localized brain structure or connectionist model can fully account for the complexity of this phenomenological experience. Prior models of flow have emphasized mechanisms such as prefrontal deactivation (Dietrich, 2004), synchronization of attentional and reward networks (Weber et al., 2009), or cerebellar contributions to internal models of motor control (Gold & Ciorciari, 2021). Our results, however, show that these accounts capture only fragments of the full picture. By applying complexity metrics to whole-brain dynamics, we reveal that flow involves a distributed, metastable coordination of neural processes that cannot be reduced to static network configurations or regional activity patterns. This suggests that existing models might benefit from integration under a broader framework that explicitly incorporates the nonlinear, emergent properties of the brain as a complex system.

The complexity science perspective also offers a way to reconcile apparent contradictions across prior studies of flow. For example, while some studies have emphasized localized hypo-frontality as central to flow, our data indicate that prefrontal contributions fluctuate dynamically depending on the dominant coordination mode. Similarly, looking beyond analysis of between-condition means to relationships between self-reported phenomenology scores and complexity metrics reveals interesting insights that go beyond simple mean differences. In this way, complexity metrics provide a unifying lens that can accommodate the diversity of empirical findings on flow.

Finally, our work highlights the potential of complexity science to advance a more neurophenomenological account of flow—one that directly links subjective experience with the dynamic interplay of large-scale brain systems. By capturing the temporal evolution of neural coordination patterns, complexity metrics move us closer to characterizing how the felt sense of effortlessness, absorption, and intrinsic reward in flow arises from underlying brain dynamics. This opens the door for future studies to more precisely map individual variations in flow experience to distinct patterns of brain organization and to explore how these dynamics shift across different contexts and forms of expertise. Ultimately, such an approach can help bridge gaps between theory, phenomenology, and neuroscience in the study of flow.

### 4.4 Limitations

The results reported within this manuscript represent a novel analysis of a pre-existing dataset (Huskey et al., 2022). Limitations associated with the dataset and experimental procedure are already well-discussed in that previous publication. We have also noted additional limitations in various subsections of this manuscript’s discussion section. Nevertheless, our new analyses introduce new limitations deserving of consideration. Notably, the myriad of complexity signatures in the neuroimaging literature may actually be limiting for their use. Many are correlated, and as demonstrated here, not all are sensitive to phenomenological or behavioral experiences. However, different signatures have been shown to be sensitive to aging populations, neo-natals, psychiatric conditions, and neurodegenerative disorders (see Hancock et al., 2025 for a review). Although it is relatively easy to estimate the signatures when, as here, the code is provided, it may be more challenging to conceptually understand the signatures and interpret the results appropriately. Moreover, we examined a vast array of complexity signatures in a data driven (rather than a priori) way. Although we took careful analytical steps to minimize the chance of false positive results, it is still important to understand these as exploratory findings requiring subsequent confirmation in a new, preferably preregistered, project (Dienlin et al., 2021).

An additional limitation concerns the interpretation of our within-condition brain-phenomenology analyses. We operationalized flow using the Autotelic Experience subscale of the Event Experience Scale (Jackson & Eklund, 2004). This subscale measures the intrinsically rewarding quality of a flow experience. At the between-condition level, the balanced-difficulty manipulation reliably elicited higher scores than the boredom or frustration conditions, a finding consistent with a successful experimental manipulation. However, when we turn to within-condition associations between complexity signatures and autotelic experience scores, the interpretation becomes more complicated. Within a condition, variation on the autotelic subscale may reflect variation in flow specifically, variation in how rewarding or pleasurable participants found the task more generally, or some combination of the two. Our data cannot adjudicate between these possibilities. Future work may use measures that more cleanly separate the multidimensional characteristics of flow (i.e., the complete Event Experience Scale; Jackson & Eklund, 2004) in order to better adjudicate this issue. This would allow stronger inferences about whether complexity signatures track flow specifically or autotelic experience more narrowly.

Finally, our analysis relies on the AAL anatomical parcellation to define regions of interest. This is a deviation from prior work on this dataset that has used an atlas defined by Seitzman and colleagues (2020). This decision was made to allow for more direct comparisons with other work using measures drawn from complexity science (e.g., Hancock, Cabral, et al., 2022; Hancock et al., 2023), although this makes comparison with prior flow research more difficult. To bridge this gap, we worked to landmark commonalities between atlases. We also note that, at a global level, measures of synchrony and metastability yield common results for both the AAL and Seitzman atlas. Nevertheless, prior work shows that some network measures are sensitive to atlas (Craddock et al., 2012; Fisher et al., 2021), and therefore future work might seek to examine these results across a multitude of atlases (Kong et al., 2025), being sure to use those that include cortical, subcortical, and cerebellar nodes (as is true for both the AAL and Seitzman atlases).

## 5. Conclusion: Flow, Media, and Wellbeing

Flow is an intrinsically rewarding and attentionally absorbing phenomenological experience that is linked with a wide variety of wellbeing outcomes (Csikszentmihalyi, 1990). In a TED talk that has attracted more than 7.8 million views, Csikszentmihalyi calls flow “the secret to happiness” (Csikszentmihalyi, 2004). A vast array of daily activities can elicit flow, including media use (Sherry, 2004), although not all media are likely to elicit flow. Low-challenge and passive media (e.g., TV viewing) rarely elicit flow, whereas as interactive media that elicit high challenge and demand high levels of player skill (e.g., video games) are much more likely to elicit flow and corresponding increases in wellbeing (for an extended discussion, see Kubey & Csikszentmihályi, 1990). Our study uses a video game to examine brain dynamics during flow. In doing so, we observe a complex system characterized by efficient information dynamics (entropy rate). Critically, these brain dynamics also explain variance in flow scores. Accordingly, we gain new insights into the ways media elicit flow, brain dynamics during flow, and linkages with wellbeing outcomes.

At the same time, we consider our work in light of high-quality evidence demonstrating that flow can be elicited from low-difficulty tasks (Melnikoff et al., 2022, 2025). This substantially contradicts long-standing theorizing that a balance between task-difficulty and individual skill is a causal precursor of flow (Csikszentmihalyi, 1990)—a theoretical framework that motivated our study design. These findings raise important questions. If flow can be elicited during low-difficulty tasks, is it flow, or more active (compared to passive) and difficult (compared to easy) media that are required for wellbeing outcomes? Considering recent findings demonstrating that high-effort tasks are perceived as valuable (Inzlicht et al., 2018) and elicit increases in wellbeing (Campbell et al., 2025), alongside uncertainty in the causal pathway between task→flow→wellbeing outcomes (Huskey & Schmälzle, 2025), it is unclear if it is flow or effort during media use that elicits wellbeing. Answering this question will help explain results showing that time (overall) spent playing video games is associated with either no (Vuorre et al., 2022) or modest increases in wellbeing (Johannes et al., 2021). Ultimately, clarifying these relationships will require careful experimentation capable of elucidating explanation at multiple levels (Huskey et al., 2020; Schmälzle & Huskey, 2023b).

Finally, there are substantial concerns about media, particularly video games, and linkages with video gaming disorder, a behavioral addiction (Weber et al., 2016; Woodman & Weber, 2025). Some evidence suggests that experiencing flow while playing video games may be a precursor to gaming disorder (Hull et al., 2013; Tokunaga, 2013), although the evidence for this claim is fairly limited (Wang et al., 2023; Weber et al., 2016). Regrettably, our study offers limited insight into this as our design is not longitudinal, and only includes healthy participants. Although, preliminary evidence demonstrates clear differences in complexity signatures between healthy and addicted video game players during passive (resting state) and active (working memory) tasks (Hosseini et al., 2021). Therefore, it seems likely that individuals experiencing gaming disorder would show different complexity signatures than the ones we observed. Fully addressing this question remains an important direction for future research.

## 6. Author Contributions

**Fran Hancock:** Conceptualization; Formal analysis; Methodology; Software; Visualization; Writing - original draft; Writing - review and editing

**Rachael Kee:** Writing - review and editing

**Fernando Rosas:** Writing - review and editing

**Manesh Girn:** Conceptualization; Writing - review and editing

**Steven Kotler:** Writing - review and editing

**Michael Mannino:** Conceptualization; Writing - review and editing

**Richard Huskey:** Conceptualization, Data Curation, Investigation, Writing - original draft, Writing - review and editing

All authors participated in the discussion of the ideas, read and approved the submitted version.

## 6.1 Funding

This research did not receive any specific grant from funding agencies in the public, commercial, or not-for-profit sectors.

## Footnotes

1 This team implicates other brain activation patterns in flow (Ulrich et al., 2016b), and many of these brain activation patterns replicate (Ulrich et al., 2022).

2 This theory draws heavily on ideas from complexity science, but constrains these ideas to specific networks.

3 We remind readers that the current manuscript re-analyzes a dataset previously reported in Huskey et al., (2022). That project also evaluated global signatures of synchrony and metastability using a slightly different data cleaning technique and analytical pipeline. Nevertheless, global synchrony and metastability results reported in that manuscript (i.e., Table 3 in Huskey et al., 2022) are largely aligned with the findings reported in this manuscript (Supplementary Figure SF7). We take this as evidence that this (null) result at the global level is not sensitive to analytical approach.

## Notes

https://osf.io/f73qm/

https://openneuro.org/datasets/ds003358/versions/1.0.0

https://osf.io/bxvhr/

https://github.com/asteroidimpact

https://github.com/franhancock/COMPLEXITY_GLOBAL_AND_LOCAL

### Competing Interest Statement

The authors have declared no competing interest.

### Summary of Updates

A revised analysis investigating the relationship between self-reported flow and entropy rate has been conduced. This resulted in a revised Figure 5. Additional text has been added to the limitations section.

